# Electrocorticography based monitoring of anaesthetic depth in mice

**DOI:** 10.1101/2021.07.12.452032

**Authors:** Dominik Schmidt, Gwendolyn English, Thomas Gent, Mehmet Fatih Yanik, Wolfger von der Behrens

## Abstract

To improve animal welfare and data quality and reproducibility during research conducted under anaesthesia, anaesthetic depth in laboratory animals must be precisely monitored and controlled. While a variety of methods have been developed to estimate the depth of anaesthesia in humans, such tools for monitoring anaesthetic depth in laboratory animals remain limited. Here we propose an epidural electrocorticogram-based monitoring system that accurately tracks the depth of anesthesia in mice receiving inhalable isoflurane anaesthesia. Several features of the electrocorticogram signals exhibit robust modulation by the concentration of the administered anesthetic, notably, corticocortical coherence serves as an excellent indicator of anaesthetic depth. We developed a gradient boosting regressor framework that utilizes the extracted features to accurately estimate the depth of anaesthesia. Our method for feature extraction and estimation is conducted with a latency of only ten seconds, establishing a system for the real-time tracking of anaesthetic depth in mice.

## 1 Introduction

General anaesthesia is commonly used in surgical procedures and acute experiments performed on laboratory animals in both fundamental and biomedical research. Exposure to general anaesthetic agents strongly perturbs multiple brain networks and can have profound, lasting effects on the physiology of exposed animals^1–3^. In order to minimize both the acute and chronic effects of anaesthesia while also safeguarding the welfare of laboratory animals during surgery, the exposure to anaesthetic agents should be expertly balanced. Namely, the administered anaesthesia should be sufficiently strong to maintain the animal in an unconscious state, while still minimally dosed to reduce the anaesthetic’s acute effect on brain function and its longitudinal effect on general physiology.

These anaesthetic constraints are well known in human practice where, to prevent post-operative complications, general anaesthesia should be titrated to avoid detrimental physiological effects^4, 5^. To facilitate an anaesthetic delivery that balances the demands of interoperative awareness and adverse effects, a significant amount of research has focused on measuring the human depth of anaesthesia (DoA)^6–8^. Such work has prominently led to the development of the proprietary Bispectral Index Score (BIS), which makes use of several electroencephalographic parameters to estimate DoA, and has been established as the predominant anaesthetic monitor used during human surgeries^4^. Other published approaches for human depth of anaesthesia estimation rely on non-linear features extracted from electroencephalographic measurements or evoked potentials^9^, which are used as inputs to traditional machine-learning algorithms or artificial neural networks^10, 11^. Other studies have further investigated using DoA measures in closed-loop control circuits to altogether replace most interventions by anaesthesiologists^4, 12^.

While methods to estimate DoA in humans are well developed and validated, for laboratory animals, and in particular mice, research and techniques remain sparse. Previous work has investigated closed-loop anaesthetic delivery in rats to control the electroencephalogram-determined burst suppression, a signature of inactivated brain states^13, 14^. Another study has linked human DoA techniques to viable methods in neonatal mice using intracortical electrophysiology^15^. Despite prior work, a gap remains in the understanding of DoA monitoring in adult mice and of the specific features of electroencephalographic signals beyond burst suppression that are modulated by anaesthesia. Further research has investigated alternative physiological measures for their usefulness in monitoring anaesthetic depth, however heart rate and blood pressure have been shown to less accurately assess DoA than the bispectral index, suggesting that methods for monitoring anaesthetic depth in mice based upon electroencephalographic signals may be most effective^16^.

Due to the limitations of the available methods to monitor DoA in small laboratory animals, we developed an epidural DoA measurement technique specifically targeted for mice. Our system achieves anaesthetic depth monitoring using inexpensive recording devices coupled with algorithms tailored to mice, the most commonly used animal model in biomedical research^17^. Similar to the human BIS, our approach makes use of electrocorticogram (ECoG) data and standard feature extraction for estimating a definition of DoA that is based on population statistics of the anaesthetic effect. Specifically, we perform estimation of the DoA on two proxies: the amplitude of somatosensory evoked potentials and the administered concentration of isoflurane. The latter yields an estimator whose momentary values are reflective of an effective isoflurane concentration. This effective isoflurane concentration is different from the administered concentration, since it is based on features directly affected by the true depth of anaesthesia, i.e. it is the administered isoflurane concentration the average mouse requires such that the ECoG signals exhibit the measured feature characteristics. Our system focuses on the application of inhalable isoflurane anaesthesia, as it allows maximum temporal control of anaesthetic depth over other injectable alternatives^18^.

Our deployed monitoring technique proves accurate in tracking the DoA across all tested animals, without the need for custom tuning to individual subjects. Feature importance analyses also suggest that corticocortical coherence is a crucial indicator of anaesthetic depth, namely that corticocortical coherence increases with increasing isoflurane concentration. Together, our results demonstrate a more robust anaesthesia monitoring system for mice and identify electrocorticogram signal features that may be useful in DoA monitoring also in other laboratory animals.

## 2 Results

Acute experiments were conducted on adult female mice. Two epidural electrocorticogram (ECoG) electrodes were placed above the right- and left-hemisphere somatosensory cortices (see Methods). Throughout the experiment, stimulation of the right whisker pad was used to evoke somatosensory responses. Concurrent to the ECoG recordings and somatosensory stimulation, animals were subjected to an anaesthesia protocol that varied the concentration of inhaled isoflurane in consecutive fifteen minute blocks (Fig. 1A and 1B). Signal features were extracted from both ECoG channels and ultimately provided as input to a gradient boosting regressor trained to estimate two proxies of anaesthetic depth, namely the administered isoflurane concentration and the sensory evoked response amplitude. Training an estimator on administered isoflurane concentration over a population of mice results in estimation of an effective isoflurane concentration, which approximates a true depth of anaesthesia. The regressor was trained and evaluated across data collected from all experimental animals in a leave-one-out cross-validation scheme, showing good generalization errors across the entire population.

**Figure 1.**
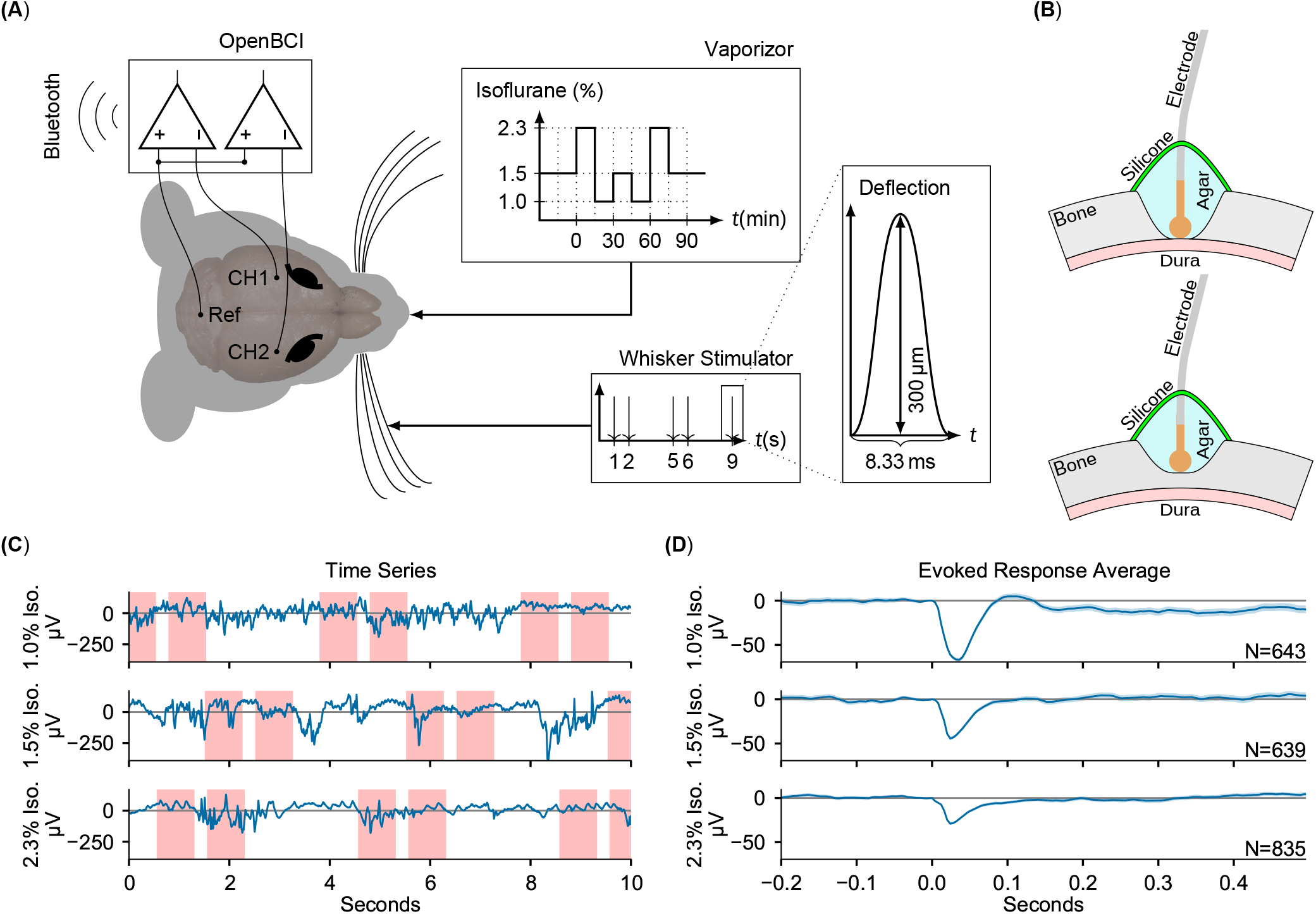
Electrocorticogram signals are extracted for estimation of anaesthetic depth. (A) Schematic overview of the recording setup. Two craniotomies over the somatosensory cortices (CH1 and CH2) were measured against a common reference over cerebellum (Ref), while stimulating the right whiskers and varying the isoflurane concentration. (B) (upper panel) Craniotomy over barrel cortices. The Ag/AgCl electrodes were placed directly on the dura, then covered in phosphate-buffered saline based agar and two component silicone. (lower panel) Partial craniotomy over cerebellum for the reference electrode, drilled to approximately 20% thickness, and covered as above. (C) Example ECoG traces recorded using the OpenBCI during different isoflurane concentrations. Red shaded areas denote the [−0.2 s,0.5 s] interval around a stimulus. (D) The evoked responses averaged over all trials in each isoflurane concentration block, zero-aligned at *t* = 0. The shaded area indicates the 2σ-range of the standard error of the mean.

### 2.1 Quality of Extracted Features

Waveform features were extracted from the ECoG signals and the effects of varying the administered isoflurane concentration were evaluated (Fig 1C-E). An exhaustive list of the median feature values across all administered isoflurane concentrations as well as the p-values of all two-sided Mann-Whitney-U-tests performed, representing the extent to which each feature exhibited anaesthetic modulation, can be found in Supplementary Tables 1 and 2. Similar to results in humans^19–21^, several features showed significant dependency on the inhaled isoflurane concentration, specifically: burst suppression ratio (*p* = 9.04 × 10^−3^), corticocortical coherence (*p* = 4.08 × 10^−5^), Lempel-Ziv complexity (*p* = 1.93 × 10^−3^), and sample entropy (*p* = 5.35 × 10^−5^) as shown in Figure 2.

**Figure 2.**
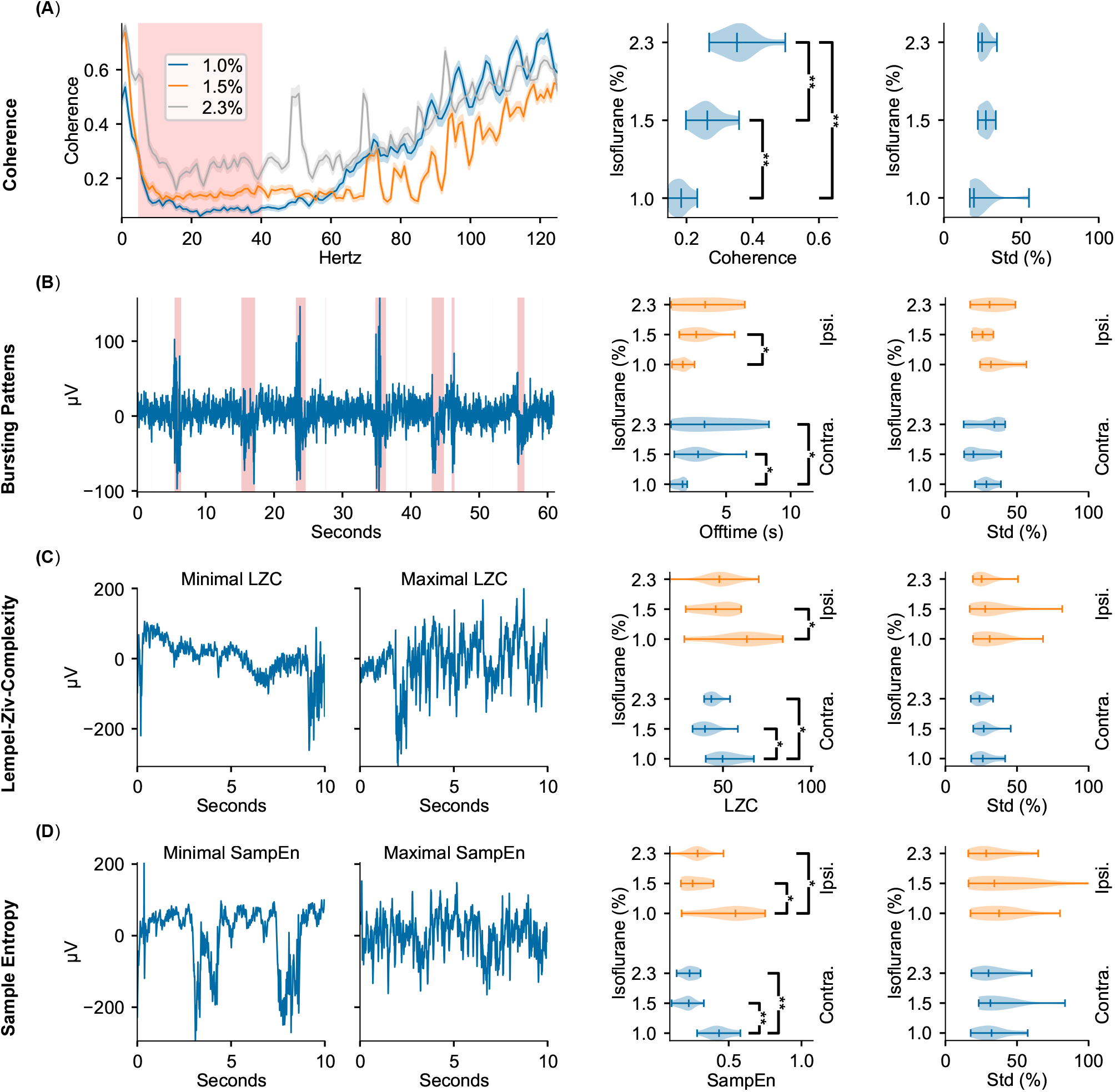
ECoG signal features display modulation to delivered isoflurane concentrations. The leftmost column depicts representative examples of the features found to be most modulated by changes in administered isoflurane concentrations. The middle column depicts violin plots of the respective feature averages over animals across isoflurane concentrations. The rightmost column depicts violin plots of the respective standard deviation within the different isoflurane segments. All violin plots display the minimum, maximum, and median values of the distributions. Row (A) Corticocortical Coherence. Representative example illustrates averages and standard error of the mean calculated over all isoflurane blocks. Highlighted region depicts the frequency range used in further analysis. Row (B) Bursting Patterns. Shaded regions in representative example depict signal portions classified as burst. Row (C) Lempel-Ziv Complexity. Representative examples of signal segments with maximum and minimum LZC. Row (D) Sample Entropy. Representative examples of signal segments with maximum and minimum sample entropy. Significance testing using two-sided Mann-Whitney U test across isoflurane concentrations. Three Benjamini-Hochberg False Detection Rate controls^23^ with decreasing false detection probability have been performed. Tests deemed significant for a false detection probability of 5% are marked with one star, for a probability of 1% with two stars and for a probability of 0.1% with three stars. Precise *p*-values are listed in supplementary Table S1.

The corticocortical spectral coherence between the two somatosensory cortex channels within the spectrum of 5 Hz to 40 Hz, comprising the theta, alpha, beta, and low gamma bands, increases with administered isoflurane concentration (see Fig. 2A). Important for DoA estimation, the corticocortical coherence exhibits highly significant differences across all tested isoflurane regimes. As depicted in Fig. 2B, increasing administered isoflurane concentration affects the burst suppression ratio by inducing more extended off (i.e. suppression) times. Increased offtime corresponding to increased administered isoflurane concentrations can be observed in both the contra- and ipsilateral hemispheres, though the former shows a higher statistical significance (*p* = 4.31*e* – 3). Notably, while the hemisphere contralateral to stimulation exhibits a significant dynamic between offtime and administered anaesthetic concentration, the on (i.e. bursting) times are not significantly effected by the administered anaesthetic concentration (see Supp Table 1), indicating that the offtime measurement may be a more effective indicator of DoA than the overall burst suppression ratio of on-to off-times.

Both Lempel-Ziv complexity (LZC, i.e. the compressibility of the signal) and sample entropy values (i.e. the randomness of the signal) peak for the lowest isoflurane concentration of 1.0%, and are significantly different from the values observed at the higher concentrations of 1.5% and 2.3% (*p* ≤ 1.28 × 10^−2^ for LZC and *p* ≤ 2.91 × 10^−3^ for sample entropy, illustrated in Fig. 2C). The increase of almost 100% for sample entropy and 20% to 30% for LZC at lower anaesthetic levels is observed in the electrode channels both contralateral and ipsilateral to the location of whisker stimulation (Fig. 2C).

The distribution of all four of these features are appreciably separated across different isoflurane concentrations, rendering them useful measures for robust estimation of anaesthetic depth. While the aforementioned features displayed consistent modulation by the administered isoflurane concentration, others we tested did not reliably vary with the anaesthesia protocol (Supplementary Tables S1 and S2). Aligned with prior research in rats^22^, modulation of the spectral edge frequency and 1/*f*-slope was not detected in our population of mice. We observed that the absolute power density reduces with increasing anaesthesia, but is subject to large inter-individual variations (a standard deviation larger than 62% of the mean), likely rendering these frequency related features ineffective in identifying consistent trends across animals without prior baseline knowledge.

Finally, we tested the features for hysteresis, which we use here to denote the amount the previous isoflurane concentration block modulates the extracted features of the current isoflurane concentration block, excluding the first five minutes after isoflurane concentration changes occur, to avoid signal transients until a steady-state is achieved (see Methods Eq. 1). A high hysteresis value denotes that the respective feature was higher on average when succeeding 2.3% isoflurane versus 1.0% isoflurane. As depicted in Fig. 3, a number of features indeed display significant hysteresis. While many features measured on the ECoG channel ipsilateral to somatosensory stimulation displayed hysteresis, this trend was not observed for features measured from the ECoG channel contralateral to stimulation. This is likely due to the large contralateral evoked responses masking the minute changes caused by the hysteresis effects between adjacent isoflurane blocks.

**Figure 3.**
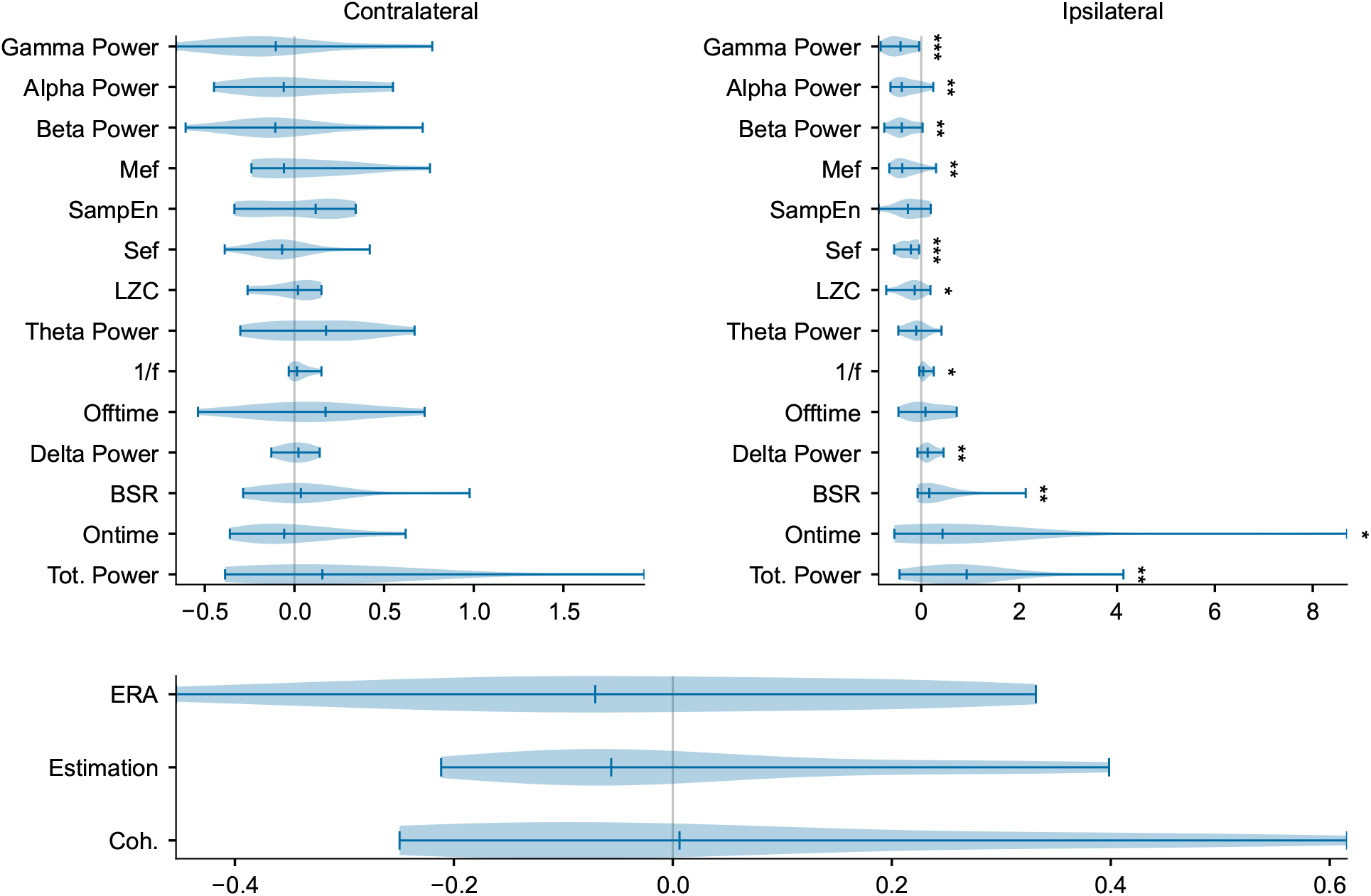
Hysteresis values indicate that anaesthetic depth estimation is not biased by previous isoflurane exposure and reveal hemispheric differences in exposure dependence. Depicted values represent the extent to which individual features are effected by the history of isoflurane exposure (see Methods Eq. 1) Statistical significance was determined with a Wilcoxon signed rank test, testing whether the median is significantly different from zero, i.e. whether the feature shows significant hysteresis. *p* < 0.05 is denoted with one asterisk, *p* < 0.01 with two and *p* < 0.001 with three. An exhaustive list of the numeric values of all p-values can be found in Supplementary Table 4

Additionally, the estimation of the isoflurane concentration and the sensory evoked response amplitude were tested for hysteresis. Neither estimation mode exhibits significant hysteresis, which can be explained by the estimator having been trained to be agnostic to any preceding isoflurane concentration.

### 2.2 Estimator Performance

The gradient boosting regressor with 100 boosting steps and maximum tree-depth of three was trained with the values of all of the previously described extracted features from the three most recent ten second recording windows (t-0,t-1,t-2) as inputs and targeted two measures of anaesthetic depth, both administered isoflurane concentration and evoked response amplitude (depicted in Fig. 4A). Gradient boosting was chosen in a pre-trial review among Support Vector Regression (SVR) with Gaussian kernels, SVR with linear kernels, K-nearest neighbors and standard linear regression after exhibiting the best performance. The estimator was trained eleven times, using a leave-one-out cross-validation scheme (i.e. for each iteration, one animal was removed from the whole data set before training and the estimation performance evaluated against that animal). Individual feature importances (measured by the Gini gain, i.e. the total estimation improvement achieved by inclusion of the feature) were then extracted and their statistics over all 11 folds were considered. Stable estimation should yield similar feature importances over all folds. Such stability was observed when estimating the effective isoflurane concentration (i.e. the estimator trained on administered concentration), however, estimation across folds was less reliable when the regressor was trained on evoked response amplitude, as shown in Fig. 4D and 4G.

**Figure 4.**
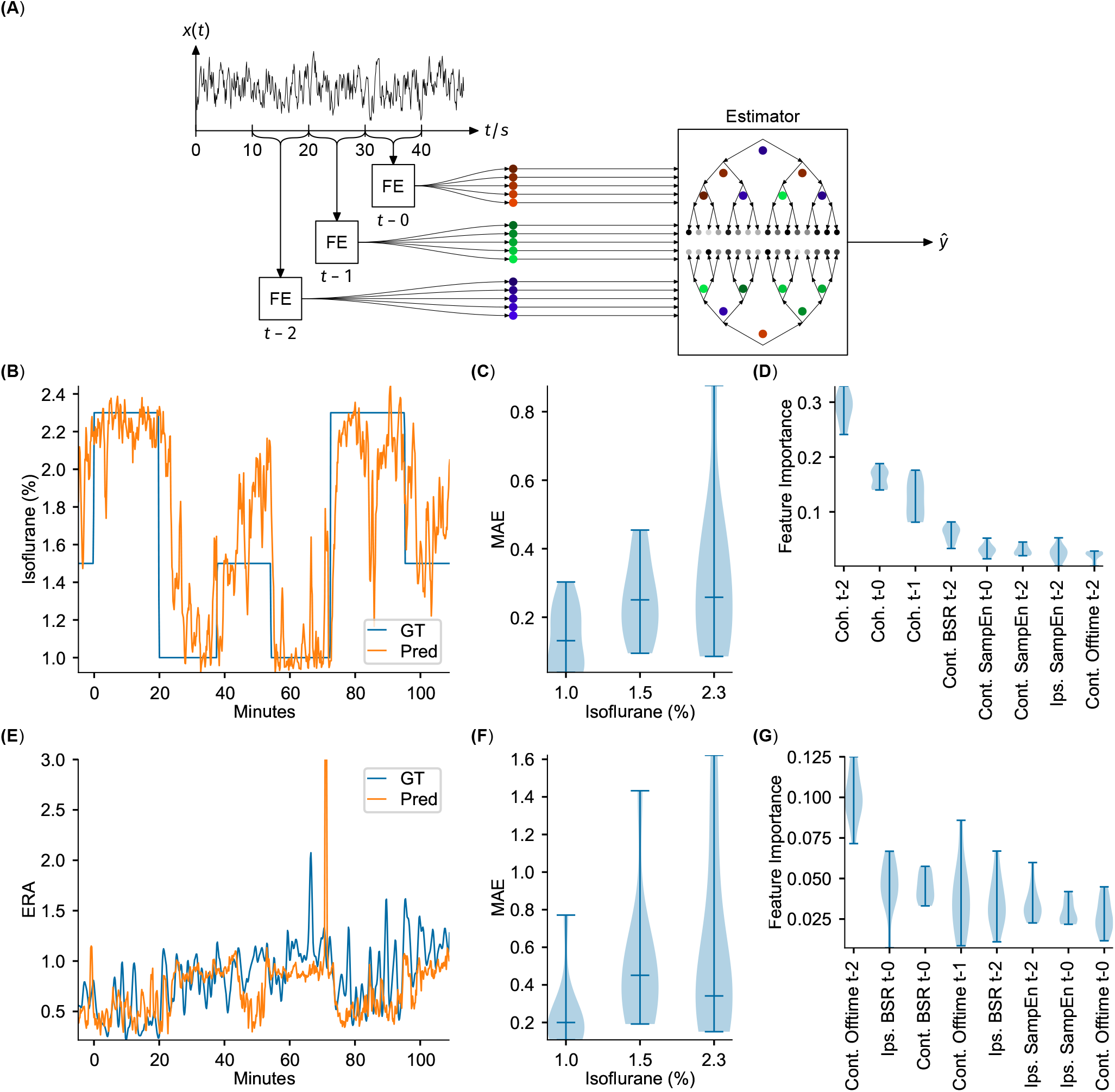
ECoG features serve as input to a gradient boosting regressor to successfully estimate DoA. Feature importances indicate corticocortical coherence as a critical DoA readout. (A) Overview of feature extraction and estimator workflow. Notch and high-pass filtered signal traces were extracted in 10 s blocks, signal features were calculated (FE), and then the three most recent consecutive blocks provided as input to estimate a target variable *ŷ* via a gradient boosting ensemble. (B) Example of the estimation of isoflurane concentration. GT denotes the measured ground-truth. (C) Distribution of the mean of the mean absolute estimation error over all animals in each isoflurane regime, corresponding to estimation of the isoflurane concentration. (D) Feature importance over all folds. The corticocortical coherences are the most important features, over almost all folds, corresponding to estimation of the isoflurane concentration. (E) Example estimation of the evoked response attenuation (ERA). GT denotes the measured ground-truth. (F) Mean absolute error over all folds or animals (*n* = 11), corresponding to estimation of the ERA. (G) Distribution of the feature importances over all animals, corresponding to estimation of the ERA. All the violin plots indicate the minimum, maximum and median values.

Estimation of the effective isoflurane concentration performed robustly over all animals (mean absolute error over folds: 0.238 ± 0.076 (mean ± std), *R*^2^-score: 0.576 ± 0.289). Fig. 4B depicts the regression results over time for an example animal and demonstrates that the estimation quickly captures changes in administered isoflurane concentration. The stability of estimating this variable is further demonstrated by the low variance in estimation performance over all 11 folds, particularly for administered isoflurane concentrations of 1.0% and 1.5%, depicted in Fig. 4C. Fig. 4D displays the feature importance extracted from all 11 folds. Coherence between the two barrel cortex electrodes proves to be critical for estimation, with the corticocortical coherence from the oldest block (i.e. *t* – 2, representing the window 30 – 20 seconds before current time, from Fig. 4D) exhibiting the strongest importance for estimation. Sample entropy in the barrel cortex electrode located ipsilateral to the whisker stimulation was also important for estimation, again with the oldest values (*t* – 2) displaying highest importance. Finally, for estimation of effective isoflurane concentration, the burst suppression ratio for the barrel cortex channel contralateral to the whisker stimulation hemisphere demonstrated to be useful.

Since the test set target variable is binned into three discrete isoflurane values, treating the isoflurane regression results as classification via nearest-neighbor quantization additionally allows for evaluation of standard multi-class classification metrics. Averaging over all One-Vs-All pairs yields an accuracy of 0.715 ± 0.134 (mean ± std), an *F*1-score of 0.699 ± 0.156, a precision of 0.780 ± 0.088 and recall of 0.714 ± 0.134.

Training the gradient boosting regressor to estimate the Evoked Response Amplitude proved less accurate and robust (mean absolute error over folds: 0.446 ± 0.252 (mean ± std), *R*^2^-score: −1.103 ± 2.009). While the estimation was accurate for several folds, or several animals, the results did not generalize well across all folds. This irregularity can be observed in Fig. 4G, which depicts highly variable feature importance.

The hysteresis analysis demonstrated that while many individual features exhibited notable changes driven by the previously presented isoflurane block, particularly in features measured on the ipsilateral channel, regressor estimation of isoflurane concentration and Evoked Response Amplitude proved robust against such effects, accurately tracking the current estimation variable irrespective of the anaesthetic depth history. Further hysteresis results are depicted in Fig. 3.

## 3 Discussion

Here we have presented an approach that utilizes epidural ECoG monitoring to estimate depth of anaesthesia in mice. Using the data collected from two electrocorticogram channels per animal, signal features were extracted and tested for their modulation by isoflurane anaesthesia. Our results indicate that many of the important features previously identified in human models translate also to mouse models, suggesting that elements of techniques developed for monitoring of human DoA may further warrant incorporation into systems targeting laboratory animals. Our analysis confirms the modulation of corticocortical coherence^24^, burst suppression ratio^25^, Lempel-Ziv complexity, and sample entropy^19^ through isoflurane anaesthesia. However, we observed a number of features that do not appear to be significantly effected by anaesthesia. Notably, the spectral measures of 1/*f*-slope^22, 26^, spectral edge frequencies and absolute power distributions either show no modulation or high inter-animal variability, rendering them unsuitable for estimation.

Isoflurane is one of the most commonly used inhalable anaesthetic agents for laboratory animals and has an inhibitory effect on excitatory neurons via a number of molecular mechanisms^2, 27^. This reduction in excitation results in a reduced stimulation of inhibitory interneurons which later causes a depletion of endogenous inhibition^28^. Such cycles of increased and reduced inhibition are manifest in the burst suppression ratio, where dose-dependent isoflurane delivery modulates the ratio of on- and off-times in mice^29^. Our experiments also reveal a dependence between administered isoflurane concentrations and the burst suppression ratio, however the computed ratio values exhibit significant differences only between the lowest administered isoflurane concentration and the higher two concentrations (e.g. 1% vs 1.5% and 2.3%). Our data indicates that the burst suppression ratio is not a strong indicator of anaesthetic depth at higher DoA. The mechanisms of isoflurane that contribute to the burst suppression ratio similarly impact the Lempel Ziv Complexity and Sample Entropy features, where induced extended offtimes create a more stable, compressible signal. Similar to our observation in the burst suppression ratio, both of these complexity features exhibit stronger significance between 1% vs 1.5% and 2.3%, again yielding their values less useful in distinguishing between higher levels of anaesthetic depth.

As indicated by the feature importances in Fig. 4D, corticocortical coherence is a robust measure for estimating depth of anaesthesia, also between higher concentrations of delivered isoflurane. The variation observed in corticocortical coherence from 5-40 Hz, comprising the theta, alpha, beta, and low gamma bands, across different anaesthetic depths has, to the best of our knowledge, not been reported in recordings made from the somatosensory cortices. Previous reports in humans and rats have identified modulation of alpha coherence in recordings made from somatosensory and frontal cortices under propofol anaesthesia^30–32^. Further studies using isoflurane in rats have indicated that coherence between primary motor and visual cortices during peripheral sensory stimulation declined as delivered anaesthetic concentrations increased^33^. Our results in recordings made from both hemispheres of the somatosensory cortex may reflect that increased isoflurane administration enforces more phase-coherence between thalamocortical projections to the two sensory hemispheres^34^. While our experiments cannot reveal the precise mechanisms of this isoflurane induced coherence, the resulting effect proves to be a critical component for estimating DoA across all depths.

Using all of the aforementioned features as input to the gradient boosting regressor yielded poor performance when the estimation of the Evoked Response Amplitude (ERA) was targeted. This poor performance could be explained by the intra- and inter-animal variability observed in the somatosensory evoked response. Previous research investigating the effects of anaesthesia in mouse models identified that the average amplitudes of visual evoked potentials were not significantly affected by variations in delivered isoflurane concentration and that evoked responses exhibited large trial-by-trial variability within delivered concentration blocks^35^. Our results, coupled with these previous findings, indicate that evoked responses recorded from primary sensory cortices across sensory modalities may not be useful readouts for DoA.

The isoflurane concentration estimator performs reliably over multiple animals as shown in Fig. 4B. All of the identified features and estimators can be evaluated with a low latency of 10 s. This evaluation latency time makes the estimation system suitable for integration into closed-loop anesthetic delivery (CLAD) systems^13, 14^, which could target specific variables reflective of the depth of anesthesia. Further enhancements could be made to estimator performance via a more exhaustive exploration of possible signal features, such as the bispectral coherence^36^, or by the implementation of neural networks to replace manual feature extraction^6, 10^.

The incorporation of our method to anaesthetized-animal biomedical research can provide numerous improvements, allowing more precise monitoring and informed adjustment of the administered anaesthesia. The use of electrocorticogram data as an input can allow the system to be incorporated into ongoing experiments without necessitating invasive intracortical electrophysiology recordings or other penetrating techniques that can otherwise compromise the integrity of the brain. Additionally, our system can assist in maintaining the appropriate balance of anaesthetic potency versus efficacy and thus contribute to the ‘Principle of Refinement’ by minimizing any conscious awareness of painful procedures by laboratory animals^37^. Our method can also provide a valuable data log for the duration of any surgical procedure that would better allow researchers to document the state of the animal throughout its experimental lifetime.

Our work can be extended for suitability to a larger set of in vivo experiments with several adjustments. We validated our system using isoflurane anaesthesia, the most common and recommended modality for acute recordings and recovery procedures in experimental mice^38^. A valuable next step would be to extend this to other anaesthetic regimens, a finding which would not only allow standardization of anaesthetic depth across experiments, but also open a window into the neuronal mechanisms involved in anaesthetic induced unconsciousness^2^. Additionally, our monitoring system could be made more flexible if the electrocorticogram signals could be recorded from different sensory modalities. Namely, if auditory or visual stimulation could be instead provided as a method for measuring changes in DoA. Such flexibility would allow researchers to select the sensory stimulation and recording sites that least interfered with their existing experimental structure.

## 4 Methods

### 4.1 Animal Experiments

In this study, eleven adult female C57BL/6J mice (age 98.0± 19.3 days, weight 21.00 ± 1.83 g) supplied by Charles River Laboratories were used for acute experiments. All experimental and surgical procedures were approved by the local veterinary authorities of the Canton Zurich, Switzerland, and were carried out in accordance with the guidelines published in the European Communities Council Directive of November 24, 1986 (86/609/EEC).

Mice were briefly induced with 3% isoflurane in oxygen anaesthesia and injected with 1mg/kg Meloxicam (Boehringer Ingelheim, Ingelheim am Rhein, Germany) as analgesic. Remaining surgical procedures were completed under 2% isoflurane in oxygen supplied at a 1L/sec flow rate. Body temperature was regulated at 37 °C with a homeothermic blanket control unit (Harvard Apparatus, Holliston, Massachusetts). Animals were mounted in a stereotactic frame and the scalp was removed to expose the skull.

Three silver electrodes were then positioned onto the skull and affixed using dental cement (Dentsply Sirona, York, Pennsylvania). Electrodes were manufactured with 250 μm thin Teflon (PTFE) coated silver wire (Goodfellow Cambridge Limited, Huntingdon, England). Teflon coating was removed to expose the wire end, which was subsequently molten to a sphere (diameter approximately 500 μm) and chlorided to improve electrochemical stability^39^.

Two craniotomies over the barrel field of primary somatosensory cortex of both hemispheres (3.5mm lateral and 1.5mm caudal of bregma) were performed to expose the intact dura. A third partial craniotomy was drilled over the cerebellar region (2mm lateral and caudal of lambda) until approximately 80% of the skull was removed. All three rounded electrode tips were then placed into the craniotomies, with both barrel cortex electrodes making contact with the dura and the reference electrode contacting the thinned skull above cerebellum. Craniotomies and electrode tips were covered with a phosphate buffered saline agar (PBS) mixture (2% PBS) (Sigma-Aldrich, St. Louis, Missouri). The agar was subsequently covered with a two-component silicone (World Precision Instruments, Sarasota, Florida) to prevent drying of the electrode sites. Silicon deposits over each craniotomy were isolated from one another to avoid electrical connectivity between recording sites. The skull surface was finally rinsed with de-ionised water to prevent parallel resistances that could interfere with the individual biological signals. Electrode impedances were typically around 10 kΩ to 20 kΩ and usually increased slightly during the recordings.

Multiple whiskers were then secured around a glass capillary placed to directly contact the center of the right hemisphere whisker field. The capillary was affixed to a piezo-bending actuator (Piezo Systems, Woburn, Massachusetts) driven by a controller with a maximum output voltage of 150V (Thorlabs, Newton, New Jersey). Whisker stimulation sequences were generated using custom LabVIEW code (National Instruments, Austin, Texas), which produced an analogue waveform with a sample rate of 200 kHz and a resolution of 16 bits. The mechanical waveforms at the capillary tip were single 120 Hz raised cosines (8.3 ms duration) with an amplitude of 300μm (1.72°) and a peak velocity of 113.1mms^−1^ (648.8 °s^−1^), confirmed using a 0.1 μm resolution laser displacement sensor (Micro-Epsilon, Ortenburg, Germany). Throughout the recording, whiskers were deflected with a 1 Hz train containing 2 second stimulus-on and stimulus-off periods (Fig. 1A). Four of the eleven mice received tail pinches delivered via Hoffman clamp to produce data used in subsequent experiments. All time periods containing tail pinch stimulation were removed from the analysed data set.

Contemporaneous with the whisker stimulation, an anaesthesia protocol was applied to achieve stable conditions at a variety of isoflurane concentrations. The protocol included fifteen minute segments of isoflurane concentration delivery in the following sequence: [1.5, 2.3, 1.0, 1.5, 1.0, 2.3, 1.5]. Isoflurane concentrations were selected to span a broad range of depths while remaining below dosages of 1.5 minimum alveolar concentrations as determined in mice^40^ and represent values of inhalable isoflurane recommended for maintenance of an anaesthetized state in mice^38^.

The completed setup was then enclosed in grounded aluminum foil for improved shielding against electromagnetic influences (e.g. from the proximal piezoelectric stimulators). Recordings were made using an open-source neural data acquisition platform (OpenBCI, Brooklyn, New York) with a gain of 24x and sample rate of 250 Hz. For all subsequent analysis, the gain factor was removed from the raw data, consequently all data reported is input referred. Electrode signals and whisker stimulation onsets were communicated to the recording computer via Bluetooth connection.

### 4.2 Feature Extraction

The recording setup yielded two electrocorticogram channels obtained from the barrel cortex electrodes located both ipsilateral and contralateral to the hemisphere of whisker stimulation. Signal traces were preprocessed using a forward-backward notch filter to remove line noise at 50 Hz and a forward-backward order Butterworth high-pass filter above 0.1 Hz to eliminate signal drift. Both raw ECoG signals were extracted from each 15 minute isoflurane concentration block. The first five minutes of each block were excluded from analysis such as to ignore the transient effects of the previous concentration block. The following spectral and time domain features of interest were then extracted from consecutive, non-overlapping 10 second sequences from each signal block.

Spectral features extracted from both electrophysiology channels included the spectral edge frequency below which 95% of the spectral power was contained (SEF 95) and the 1/*f*-slope fit over the frequency range 20 Hz to 40 Hz. Additionally, the corticocortical coherence was calculated between the two electrophysiology channels and averaged over the frequency range 5 Hz to 40 Hz. Finally, spectral power content in multiple EEG bands^41^ were extracted, namely in the *δ* (0.1 Hz to 4 Hz), *θ* (4 Hz to 8 Hz), *α* (8 Hz to 13 Hz), *β* (13 Hz to 30 Hz, and *γ* (above 30 Hz) bands. All spectral features are based on Welch’s power spectral density estimation calculated independently over each 10 s window.

To complement the spectral features, a variety of time domain features were identified for further use in the algorithmic determination of anaesthetic depth. The sample entropy was computed over signal subsequences of 80 ms length. As a measure of compressibility of the electrophysiology channels, median-thresholded signals were used to compute the Lempel-Ziv complexity (LZC). The burst suppression ratio (BSR) was calculated over the entire isoflurane concentration block to quantify the ratio of periods of high and low signal activity. Finally, the evoked response amplitude (ERA) was computed by a moving-window average of the trial-by-trial maximum evoked response amplitude to whisker stimulation and comparing the averaged values across isoflurane concentrations blocks to those computed during the 1.0% isoflurane block.

The described features were tested for modulation by isoflurane through averaging them over segments of equal isoflurane concentration for each mouse. The significance of this modulation was quantified by executing a Mann-Whitney-U-Test for every pair of unequal isoflurane concentrations for each feature. To correct for multiple comparisons, a Benjamini-Hochberg False Detection Rate control^23^ has been applied. Further details on feature calculations can be found in the supplementary material.

#### 4.2.1 Hysteresis

Hysteresis values were calculated by averaging each feature *i* in both 1.5% isoflurane segments, and calculating the difference of this mean when succeeding 1.0% isoflurane 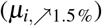 and 2.3% 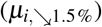 isoflurane. In order to make the different features comparable, this difference was normalized over the mean at 1.0%. The hysteresis *h_i_* of each feature *i* is thus:

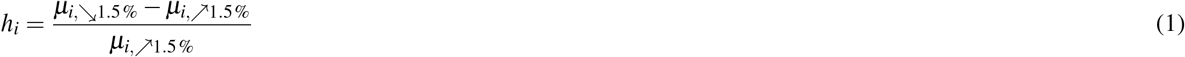

### 4.3 Estimation

As anaesthetic depth is a value on a continuous spectrum, we perform a regression analysis to identify the current anaesthetic state of the animal based on the extracted features. The features were provided as inputs to a gradient boosting regressor implemented by the python scikit-learn package^42^. Such gradient boosting algorithms combine weak estimators *g_m_* into a strong estimator *G_m_* via superposition:

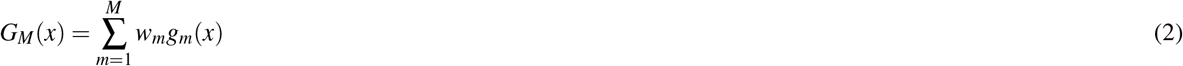

with 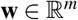 weighting the individual estimators. Here the individual estimators are chosen to be shallow decision trees (with a maximum depth of three).

A decision tree is a binary tree whose leaf nodes are assigned a real valued number – the estimation. On every non-leaf node, a decision is made what child node to traverse to, based on a single feature and a threshold for that feature. Estimation using decision trees is thus a simple tree traversal, returning the value assigned to the leaf. In the training process, the optimal feature and threshold is determined for every node by iterating through all possible features, determining the optimal threshold (which can be done computationally efficient), and greedily choosing the feature and threshold that minimizes the error^43^. The decrease in error after the split based on this feature is tracked globally, which (summed over all decision trees and normalized over all features) is represented by the Gini gain computed in feature importance calculations^44^. In gradient boost regression, every successive estimator is trained on the residual error of the superposition of all previous ones, resulting in successively improved overall loss^43^.

Two target variables for the estimators were used. First, one estimator was trained on the Evoked Response Amplitude (ERA). A second gradient boosting estimator was trained to estimate the known administered isoflurane concentration. Estimating the administered isoflurane concentration can be understood as an anaesthetic depth estimation similar to the bispectral index score. A population of mice has an average reaction to any given isoflurane concentration. Training over enough data yields an estimator that assigns the most probable isoflurane concentration at which the average mouse (over the training population) would have the reaction observed. We can further show that, with certain assumptions, estimating isoflurane concentration is equivalent to estimating the true depth of anaesthesia. We do this by modelling the DoA problem as a Bayesian network, with probability density function 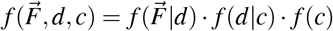 where 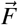 is the extracted features, *d* the true depth of anaesthesia (which is a hidden variable), and c the administered isoflurane concentration.

An optimal estimator (minimizing a square loss), which has perfect knowledge of the density 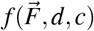, will estimate 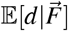^43^. Since *d* is a hidden variable, we have no way to train on it, and thus settle on estimating 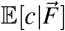. How does this surrogate compare to the optimal estimation?

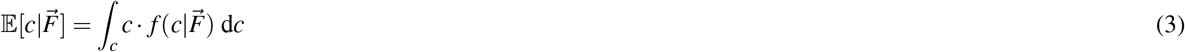

This can be expanded if we marginalize over the hidden variable *d*, and by applying the Bayes rule we get:

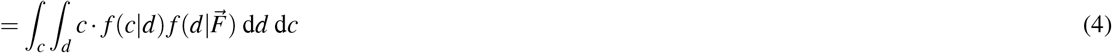

Swapping the integrals results in an inner expectation:

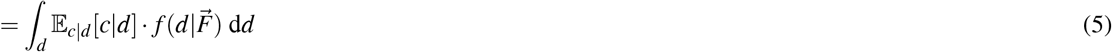

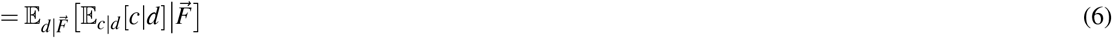

We can see that what separates this estimator from the optimal one is a transformation from anaesthetic depth to the average isoflurane concentration *h* : *d* ↦ *c*, 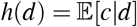. Assuming that *h* is sufficiently linear where 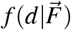 has most of its support, we can apply the linearity of the expectation and get the following approximation:

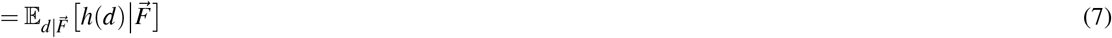

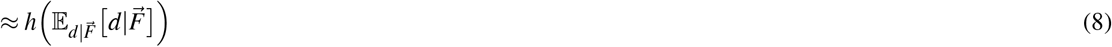

It is thus approximately a linear transformation of the optimal estimation.

While the training data on the isoflurane concentration estimation is binned into the discrete values 1.0, 1.5, 2.3, a regression approach was preferred to classification nonetheless, to capture the continuous nature of anaesthetic depth and to better gauge the promptness of response without delays due to the quantization inherent to classification.

As input, the estimator was provided the values of each feature *x* in the set of all extracted features **x** described above at three consecutive ten second sub-sequences, yielding a data set of

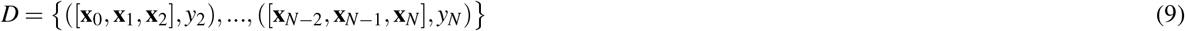

where **x**_n_ represents the feature vector at the *n*-th 10 second sub-sequence, and *y* represents the anaesthetic depth, given as either the administered isoflurane concentration of evoked response amplitude, in the same sub-sequence.

Using the data sets acquired from the *n* = 11 mice, leave-one-out cross-validation was performed, with each fold excluding the data set recorded from one animal. The estimator was trained on the remaining *n* = 10 data sets, and evaluated on the excluded test animal, yielding an estimation of the generalization error. For estimation of the administered isoflurane, a surrogate classification metric has been used, by binning the estimated isoflurane concentration into the three bins 1.0, 1.5, 2.3 via nearest-neighbor quantization and calculating the 3-class classification scores with averaging over all One-vs-All pairs.

With this approach an estimator is established, capable of making online anaesthetic depth predictions by requiring a total of only 30 seconds of feature data for estimation, with a minimal latency of 10 seconds.

## 5 Acknowledgments

This work was funded by the Swiss National Science Foundation (Project Grant Nr. 310030 172962). TG was supported by the University of Zürich Forschungskredit (FK-017-64) and the Federal Food Safety and Veterinary Office of Switzerland (2.20.02). The authors would like to thank Markus Marks for his valuable help with machine learning methods.

## 6 Author contributions statement

G.E. and W.B. conceived the study. G.E. and D.S. designed the experiments and recording techniques. D.S. conceived of and conducted all data analysis. G.E.,T.G., and D.S. performed the experiments. D.S. and G.E. wrote the manuscript with review from W.B., M.F.Y. and T.G.

