## Supplementary Material for "Electrocorticography based monitoring of anaesthetic depth in mice"

### 1 Supplementary Data

#### 1.1 Feature Extraction

For determining the burst suppression ratio, the detection of bursts is completed via simple thresholding of the high-pass filtered signal (0.1 Hz first order Butterworth forward-backward high-pass). The threshold-value $\lambda$ is calculated by taking the minimum of the local standard deviation of the signal. The local standard deviation of the signal is calculated by first low-pass filtering the high-pass filtered data $y_{HP}$ with $H$ to receive the local mean $m[n] = (H * y_{HP})[n]$, and then low-pass filtering the square distance from that mean, $\sigma^2[n] = (y_{HP}[n] - m[n])^2 * H[n]$. The threshold is hence $\lambda = \min_n \sqrt{|\sigma^2[n]|}$, with $H$ a forward-backward first order Butterworth low-pass at $1/300$ Hz. This results in the binary sequence $b[n] = y_{HP}[n] < -\lambda$. The burst suppression ratio is then calculated as the local average of this sequence, i.e. $\mathrm{BSR}[n] = b * G$ with $G$ a forward-backward first order Butterworth low-pass at $1/120$ Hz. The on- and off-times are easily extracted directly from the binary sequence. We observed that mostly peak values change with isoflurane, hence an envelope filter was applied over the on- and off-times. This *non-linear* filter is defined as:

$$E[n] = \max\left\{y[n],\ y[n]b[0] + y[n-1]b[1] - a[1]E[n-1]\right\} \tag{1}$$

With $a[0] = 1, a[1], b[0], b[1]$ the coefficients of a standard first order Butterworth low-pass filter at $1/100$ Hz. Unlike all other features, the bursting pattern is thus calculated continuously for the whole recording, and not in 10 s signal subsequences.

The *Lempel-Ziv Complexity* (LZC) is a measure of the compressibility of a signal. Each 10 second signal sequence was first thresholded against the median value, yielding a binary sequence which was then compressed using the LZC algorithm. The LZC value is then computed as the ratio of compressed to uncompressed binary sequence length[1].

The *Sample Entropy* is a measure of complexity of subsequences of a time-series signal[2]. Similar signal subsequences of length 80 ms were identified, then the signal values at the next time-step were compared to yield the sample entropy value.

The *Evoked Response Attenuation* (ERA) is extracted by evaluating the evoked responses in the interval $[0\,\mathrm{s}, 0.2\,\mathrm{s}]$, applying a moving window average (20 presentation in the past, 20 in the future) over the presentations, and extracting the minimal value in that interval. The attenuation is calculated by dividing these instantaneous values against their average under 1.0 % isoflurane.

The *Spectral Edge Frequency* is a measure of power distribution. The spectral edge frequency was calculated to identify the frequency below which 95 % of the spectral power is contained.

The $1/f$*-slope* is the $\alpha$-parameter of a curve-fit of the power spectral density to a function $\hat{S}_x x(f) = \frac{a}{f^\alpha}$. This has been done for the frequency range of 20 Hz to 40 Hz by first transforming the power spectral density into a log-log-space, and then via least-squares line fit.

The *Coherence* feature measures the correlation of two signals over frequency regions.The coherence between the two barrel cortex electrodes was calculated and the value averaged over 5 Hz to 40 Hz.

### 2 Supplementary Tables and Figures

#### 2.1 Figures

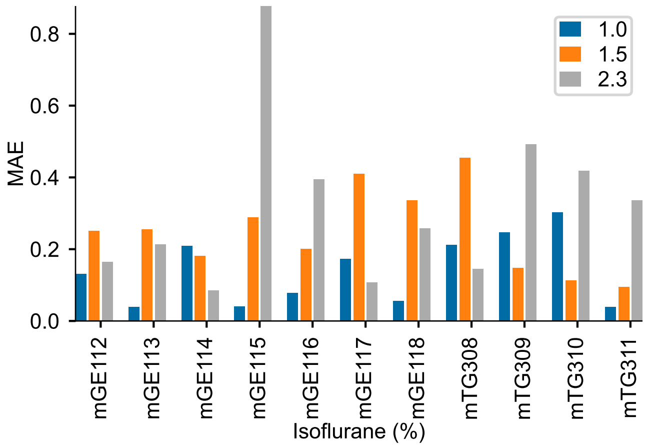


**Figure 1.** Minimum Absolute Error of Isoflurane Estimation of each mouse.

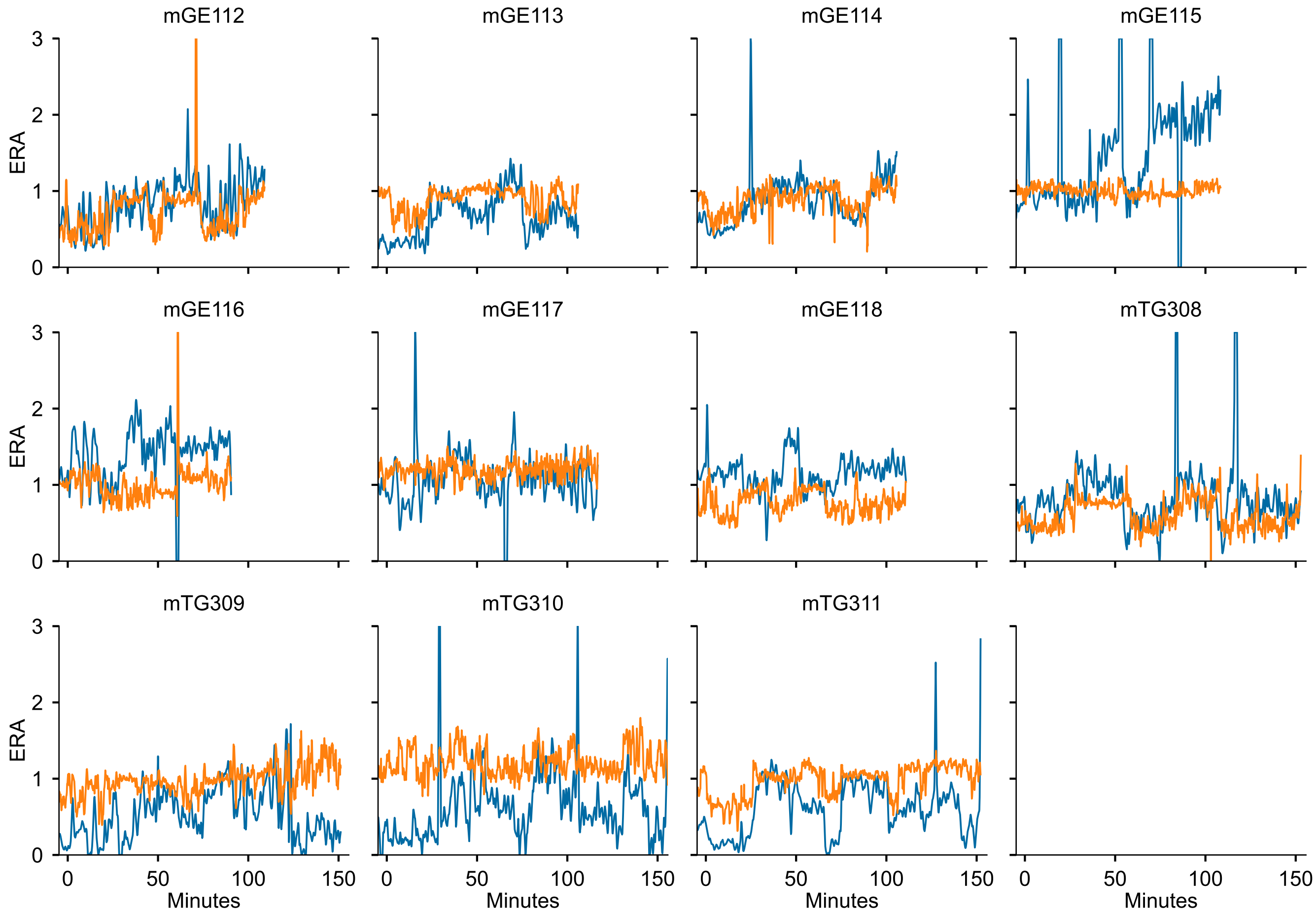


**Figure 2.** The ERA estimation results of all the 11 folds. The traces shown are always evaluated on a dataset not seen in the training data.

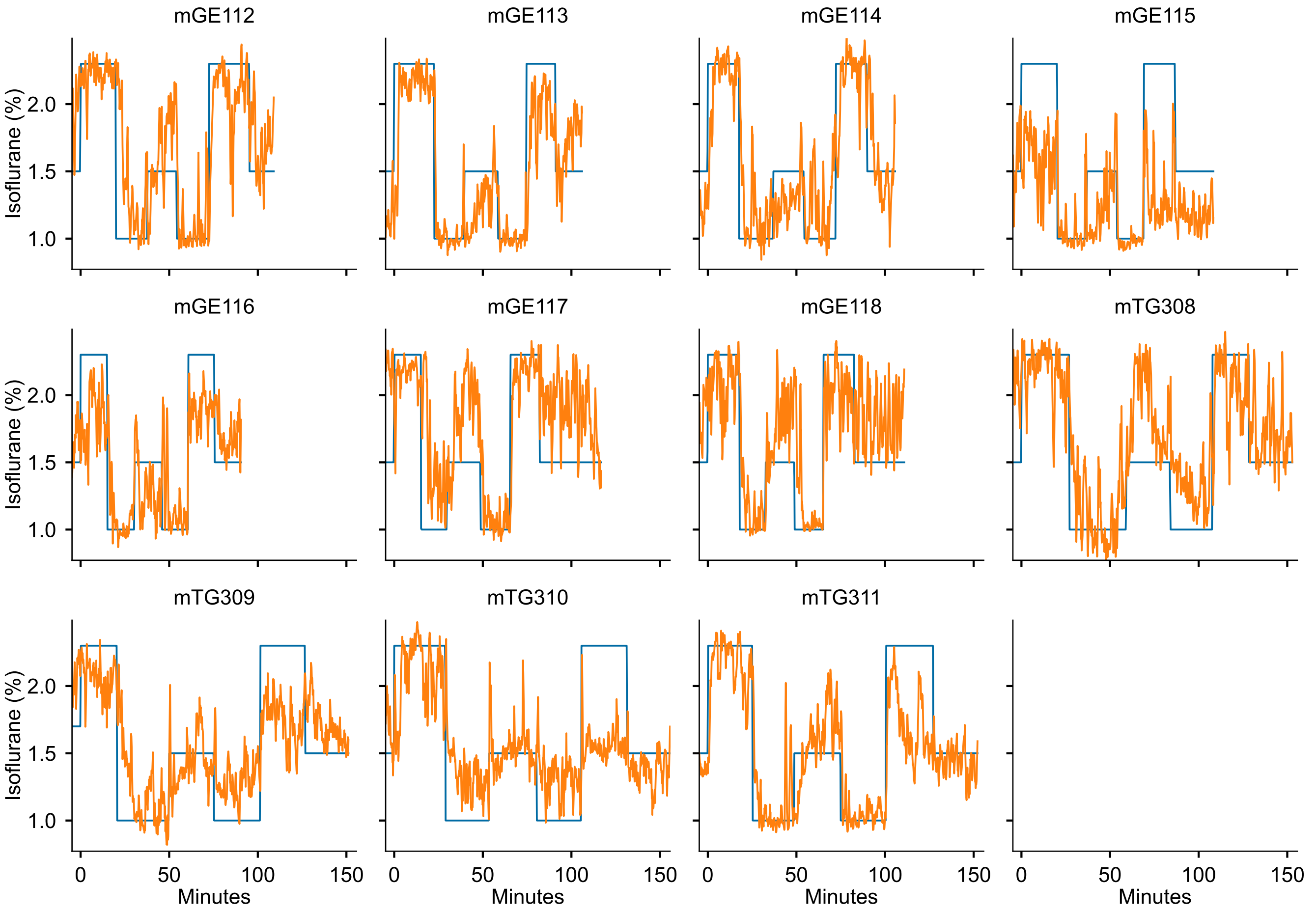


**Figure 3.** The administered isoflurane estimation results of all 11 folds. The traces shown are always evaluated on a dataset not seen in the training data.

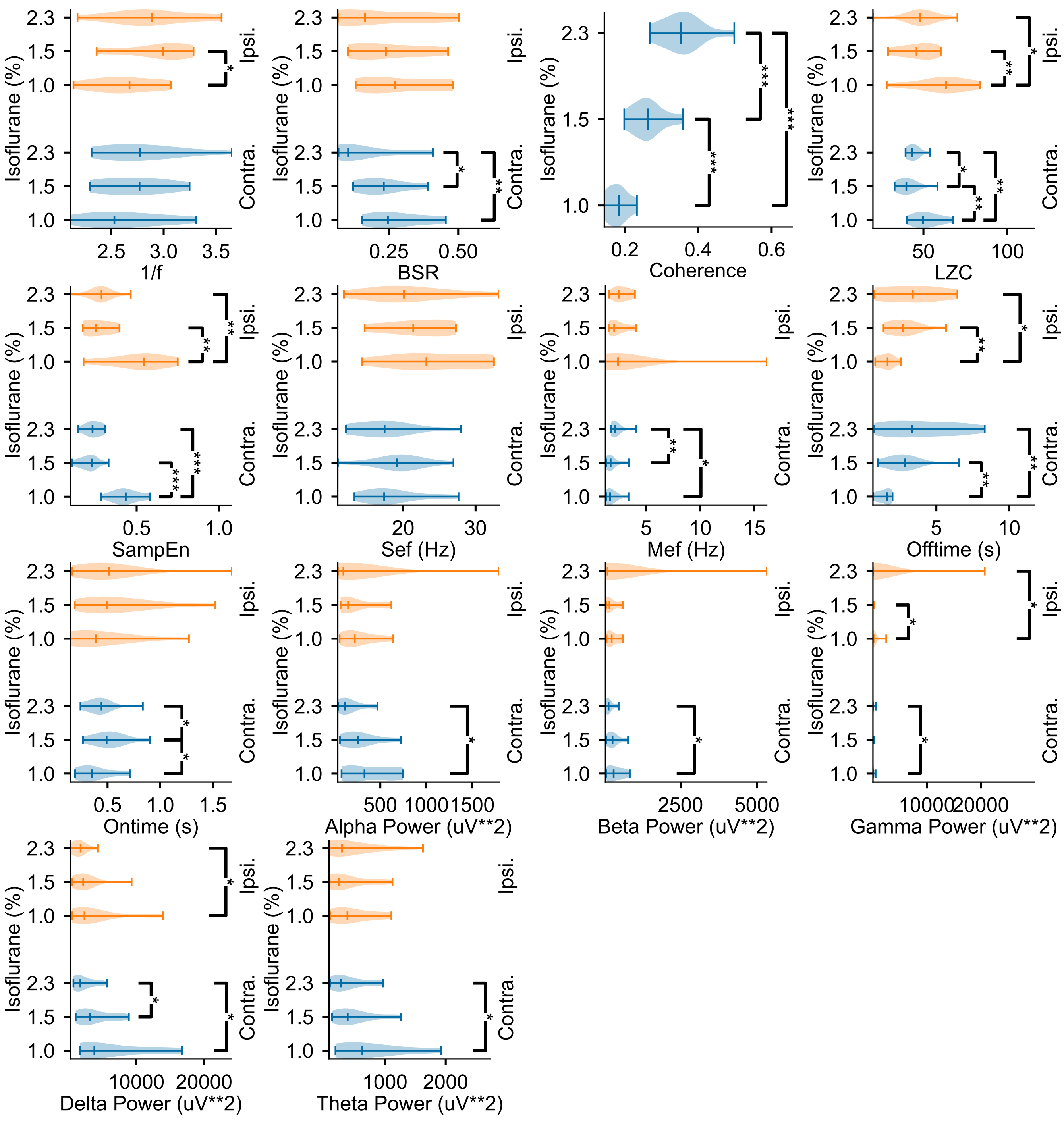

**Figure 4.** The modulation of all extraced features by isoflurane concentration

##### 2.2 Tables

| Feature | Channel<br>Isoflurane<br>Isoflurane | Ipsilat.<br>1.5 | <br>2.3 | Contralat.<br>1.5 | <br>2.3 |
|---|---|---|---|---|---|
| Coherence | 1.0 | $\mathbf{1.52 \times 10^{-4}}$ | $\mathbf{4.08 \times 10^{-5}}$ | $\mathbf{1.52 \times 10^{-4}}$ | $\mathbf{4.08 \times 10^{-5}}$ |
| | 1.5 | | $\mathbf{6.46 \times 10^{-4}}$ | | $\mathbf{6.46 \times 10^{-4}}$ |
| SampEn | 1.0 | $\mathbf{5.35 \times 10^{-5}}$ | $\mathbf{9.10 \times 10^{-5}}$ | $\mathbf{1.56 \times 10^{-3}}$ | $\mathbf{2.91 \times 10^{-3}}$ |
| | 1.5 | | $1.47 \times 10^{-1}$ | | $3.96 \times 10^{-1}$ |
| Offtime | 1.0 | $\mathbf{1.01 \times 10^{-3}}$ | $\mathbf{9.04 \times 10^{-3}}$ | $\mathbf{4.31 \times 10^{-3}}$ | $\mathbf{2.44 \times 10^{-2}}$ |
| | 1.5 | | $1.79 \times 10^{-1}$ | | $2.56 \times 10^{-1}$ |
| LZC | 1.0 | $\mathbf{1.93 \times 10^{-3}}$ | $\mathbf{4.31 \times 10^{-3}}$ | $\mathbf{6.29 \times 10^{-3}}$ | $\mathbf{1.28 \times 10^{-2}}$ |
| | 1.5 | | $\mathbf{3.81 \times 10^{-2}}$ | | $4.48 \times 10^{-1}$ |
| Mef | 1.0 | $3.96 \times 10^{-1}$ | $\mathbf{2.09 \times 10^{-2}}$ | $4.22 \times 10^{-1}$ | $1.79 \times 10^{-1}$ |
| | 1.5 | | $\mathbf{6.29 \times 10^{-3}}$ | | $1.19 \times 10^{-1}$ |
| BSR | 1.0 | $1.06 \times 10^{-1}$ | $\mathbf{9.04 \times 10^{-3}}$ | $1.62 \times 10^{-1}$ | $1.06 \times 10^{-1}$ |
| | 1.5 | | $\mathbf{3.81 \times 10^{-2}}$ | | $2.35 \times 10^{-1}$ |
| Delta Power | 1.0 | $1.97 \times 10^{-1}$ | $\mathbf{1.28 \times 10^{-2}}$ | $2.56 \times 10^{-1}$ | $\mathbf{3.81 \times 10^{-2}}$ |
| | 1.5 | | $\mathbf{2.84 \times 10^{-2}}$ | | $1.32 \times 10^{-1}$ |
| Total Power | 1.0 | $1.79 \times 10^{-1}$ | $\mathbf{1.78 \times 10^{-2}}$ | $1.97 \times 10^{-1}$ | $8.40 \times 10^{-2}$ |
| | 1.5 | | $\mathbf{3.30 \times 10^{-2}}$ | | $2.35 \times 10^{-1}$ |
| Alpha Power | 1.0 | $2.35 \times 10^{-1}$ | $\mathbf{1.78 \times 10^{-2}}$ | $3.00 \times 10^{-1}$ | $2.15 \times 10^{-1}$ |
| | 1.5 | | $9.45 \times 10^{-2}$ | | $2.77 \times 10^{-1}$ |
| Beta Power | 1.0 | $2.56 \times 10^{-1}$ | $\mathbf{1.78 \times 10^{-2}}$ | $1.79 \times 10^{-1}$ | $7.43 \times 10^{-2}$ |
| | 1.5 | | $8.40 \times 10^{-2}$ | | $2.15 \times 10^{-1}$ |
| Gamma Power | 1.0 | $1.32 \times 10^{-1}$ | $\mathbf{3.81 \times 10^{-2}}$ | $\mathbf{2.09 \times 10^{-2}}$ | $\mathbf{4.39 \times 10^{-2}}$ |
| | 1.5 | | $1.79 \times 10^{-1}$ | | $3.47 \times 10^{-1}$ |
| Ontime | 1.0 | $\mathbf{4.39 \times 10^{-2}}$ | $1.47 \times 10^{-1}$ | $1.32 \times 10^{-1}$ | $2.56 \times 10^{-1}$ |
| | 1.5 | | $\mathbf{2.44 \times 10^{-2}}$ | | $3.47 \times 10^{-1}$ |
| 1/f | 1.0 | $3.00 \times 10^{-1}$ | $2.15 \times 10^{-1}$ | $\mathbf{2.84 \times 10^{-2}}$ | $5.75 \times 10^{-2}$ |
| | 1.5 | | $3.47 \times 10^{-1}$ | | $2.77 \times 10^{-1}$ |
| Theta Power | 1.0 | $1.62 \times 10^{-1}$ | $\mathbf{3.30 \times 10^{-2}}$ | $2.35 \times 10^{-1}$ | $2.77 \times 10^{-1}$ |
| | 1.5 | | $1.79 \times 10^{-1}$ | | $5.00 \times 10^{-1}$ |
| Sef | 1.0 | $1.79 \times 10^{-1}$ | $3.23 \times 10^{-1}$ | $1.79 \times 10^{-1}$ | $9.45 \times 10^{-2}$ |
| | 1.5 | | $8.40 \times 10^{-2}$ | | $2.35 \times 10^{-1}$ |

**Table 1.** P-Values of the two-sided Mann-Whitney-U test between the feature mean distributions in different isoflurane concentrations. $n = 11$

| | 1.0 | 1.5 | 2.3 |
|---|---|---|---|
| Contralateral 1/f | 2.66 | 2.76 | 2.81 |
| Ipsilateral 1/f | 2.67 | 2.95 | 2.88 |
| Coherence | 0.17 | 0.25 | 0.35 |
| Contralateral BSR | 0.25 | 0.23 | 0.12 |
| Ipsilateral BSR | 0.26 | 0.25 | 0.13 |
| Contralateral LZC | 52.00 | 40.00 | 45.00 |
| Ipsilateral LZC | 64.00 | 46.00 | 50.00 |
| Contralateral Mef | 1.59 Hz | 1.70 Hz | 1.99 Hz |
| Ipsilateral Mef | 1.97 Hz | 1.91 Hz | 2.58 Hz |
| Contralateral Offtime | 1.40 s | 2.54 s | 3.52 s |
| Ipsilateral Offtime | 1.34 s | 2.41 s | 3.31 s |
| Contralateral Ontime | 0.32 s | 0.46 s | 0.37 s |
| Ipsilateral Ontime | 0.31 s | 0.43 s | 0.38 s |
| Contralateral SampEn | 0.42 | 0.20 | 0.23 |
| Ipsilateral SampEn | 0.53 | 0.26 | 0.29 |
| Contralateral Sef | 17.73 Hz | 19.24 Hz | 16.51 Hz |
| Ipsilateral Sef | 23.41 Hz | 21.75 Hz | 19.64 Hz |
| Contralateral Alpha Power | 232.23 $\mu V^2$ | 250.67 $\mu V^2$ | 112.27 $\mu V^2$ |
| Ipsilateral Alpha Power | 160.18 $\mu V^2$ | 159.28 $\mu V^2$ | 81.44 $\mu V^2$ |
| Contralateral Beta Power | 239.70 $\mu V^2$ | 266.35 $\mu V^2$ | 102.10 $\mu V^2$ |
| Ipsilateral Beta Power | 169.88 $\mu V^2$ | 169.94 $\mu V^2$ | 73.34 $\mu V^2$ |
| Contralateral Gamma Power | 73.15 $\mu V^2$ | 63.51 $\mu V^2$ | 25.40 $\mu V^2$ |
| Ipsilateral Gamma Power | 65.66 $\mu V^2$ | 47.92 $\mu V^2$ | 24.36 $\mu V^2$ |
| Contralateral Delta Power | 3735.47 $\mu V^2$ | 2799.92 $\mu V^2$ | 1524.08 $\mu V^2$ |
| Ipsilateral Delta Power | 2172.46 $\mu V^2$ | 1848.03 $\mu V^2$ | 960.36 $\mu V^2$ |
| Contralateral Theta Power | 466.57 $\mu V^2$ | 415.59 $\mu V^2$ | 318.52 $\mu V^2$ |
| Ipsilateral Theta Power | 328.25 $\mu V^2$ | 291.60 $\mu V^2$ | 197.21 $\mu V^2$ |

**Table 2.** Median values of all features, over all $n = 11$ mice.

| Channel<br>Feature | Ipsilat. | Contralat. |
|---|---|---|
| Sef | $1.75 \times 10^{-1}$ | $\mathbf{9.77 \times 10^{-4}}$ |
| Gamma Power | $2.40 \times 10^{-1}$ | $\mathbf{9.77 \times 10^{-4}}$ |
| Beta Power | $4.13 \times 10^{-1}$ | $\mathbf{1.95 \times 10^{-3}}$ |
| Mef | $4.13 \times 10^{-1}$ | $\mathbf{6.84 \times 10^{-3}}$ |
| Total Power | $2.40 \times 10^{-1}$ | $\mathbf{6.84 \times 10^{-3}}$ |
| Delta Power | $8.31 \times 10^{-1}$ | $\mathbf{6.84 \times 10^{-3}}$ |
| BSR | $5.77 \times 10^{-1}$ | $\mathbf{9.77 \times 10^{-3}}$ |
| Alpha Power | 1.00 | $\mathbf{9.77 \times 10^{-3}}$ |
| LZC | $8.98 \times 10^{-1}$ | $\mathbf{1.37 \times 10^{-2}}$ |
| 1/f | $1.47 \times 10^{-1}$ | $\mathbf{1.86 \times 10^{-2}}$ |
| Ontime | $7.65 \times 10^{-1}$ | $\mathbf{2.44 \times 10^{-2}}$ |
| SampEn | $8.98 \times 10^{-1}$ | $5.37 \times 10^{-2}$ |
| Theta Power | $1.75 \times 10^{-1}$ | $1.75 \times 10^{-1}$ |
| Offtime | $5.77 \times 10^{-1}$ | $4.13 \times 10^{-1}$ |
| Coherence | $8.31 \times 10^{-1}$ | $8.31 \times 10^{-1}$ |

**Table 3.** P-Values of the Wilcoxon signed-rank tests performed testing features for hysteresis.
